# Classification of early and late stage Liver Hepatocellular Carcinoma patients from their genomics and epigenomics profiles

**DOI:** 10.1101/292854

**Authors:** Harpreet Kaur, Sherry Bhalla, GPS Raghava

## Abstract

**Background:** Liver Hepatocellular Carcinoma (LIHC) is the second major cancer worldwide, responsible for millions of premature deaths every year. Prediction of clinical staging is vital to implement optimal therapeutic strategy and prognostic prediction in cancer patients. However, to date, no method has been developed for predicting stage of LIHC from genomic profile of samples.

**Results:** In current study, in silico models have been developed for classifying LIHC patients in early and late stage using RNA expression and DNA methylation data. The Cancer Genome Atlas (TCGA) dataset contains 173 early and 177 late stage samples of LIHC, was extensively analysed to identify differentially expressed RNA transcripts and methylated CpG sites that can discriminate early and late stages of LIHC samples with high precision. Naive Bayes model developed using 51 features that combine 21 CpG methylation sites and 30 RNA transcripts achieved maximum MCC 0.58 with accuracy 78.87% on validation dataset. Further, we also analysed genomics and epigenomics profiles of normal and LIHC samples and developed model to classify LIHC samples with AUROC 0.99. In addition, multiclass models developed for classifying samples in normal, early and late stage of cancer and achieved accuracy of 76.54% and AUROC of 0.86.

**Conclusion:** Our study reveals stage prediction of LIHC samples with high accuracy based on genomics and epigenomics profiling is a challenging task in comparison to classification of LIHC and normal samples. Comprehensive analysis, differentially expressed RNA transcripts, methylated CpG sites in LIHC samples and prediction models are available from CancerLSP ( http://webs.iiitd.edu.in/raghava/cancerlsp/).

## Background

Hepatocellular Carcinoma (HCC) is the fifth most common cancer and considered as the second major cause of cancer-related mortality with nearly 7,88,000 deaths occurring worldwide due to liver cancer in the year 2015. Further in United States, approximately 30,200 deaths and 42,220 new cases are estimated in 2018. It is nearly four times more frequent in males than females [1]. Also, the higher number of HCC cases is reported in Africa and Asia in comparison to Europe, these observations indicate that HCC is defined by various factors. The pathogenesis of HCC is contributed by some risk factors like viral hepatitis infection (hepatitis B or C) or cirrhosis, smoking, alcohol and lifestyle *etc* [2]. Despite improved screening and discoveries, HCC exhibits rapid clinical course with elevated mortality rate. Patients with HCC are usually identified at advanced stages due to lack of pathognomonic symptoms, which consequently limits the potential treatment options and early death [3]. Furthermore, there is high recurrence rate of 70% even after curative resection treatment [4]. Therefore, absolute cure of this disease is quite challenging, indicating an urgent need for identification of sensitive diagnostic and prognostic markers for HCC [5,6].

Traditionally in most of the developing countries, alpha-fetoprotein (AFP) is extensively employed as HCC biomarker. Its level becomes detectable in HCC once tumor is in advanced stage [7,8]. Besides, AFP-L3, a glycoform of AFP (AFP reacts with Lens culinaris agglutinin) has also been employed as HCC biomarker due to its higher sensitivity and specificity than alone AFP. The lack of reliability, insufficient sensitivity and specificity are the major associated limitations with these markers [9]. Also, Des-gamma-carboxyprothrombin (DCP) is another vital HCC biomarker. Its levels have shown to be upregulated in advanced stages [10–12]. In recent times, next-generation sequencing technology and bioinformatics analysis emergence has facilitated the identification of tumor and progression biomarkers of HCC [13]. Anomalous expression of cancer-associated genes is one of the major causes of tumorigenesis and plays vital role in hepatocarcinogenesis [14]. Evidently various reports have shown the elevated expression of *USP22, CBX6, NRAGE, ACTL6* and *CHMP4B* genes correlates with larger tumor size, advanced tumor stages, poor prognosis and short survival time of patients in HCC [15–19]. Moreover, the downregulation of *BTG1, FOXF2* and *CYP3A5* genes, have been observed to link with poorly differentiated and aggressive tumors, shorter disease survival rates and shorter recurrence times in HCC [20–22]. Beside this, recently, long non-coding RNAs have also been found to be aberrantly expressed and implicated in HCC pathogenesis [23–25]. *ZEB1- AS1* and *ANRIL* expression observed to be upregulated and their association with higher histological grade and stage in HCC [26,27]. Hence, the potential reversibility of epigenetic abnormalities and restoration of the expression of tumor suppressor genes or other genes by specific inhibitors offer a rational therapeutic approach for HCC. Though in recent past, numerous tumor drivers like *TERT, CTNNB1 etc* have been identified in HCC, however, most of them have not been translated into efficient modalities [28].

In addition, various studies have shown that the alterations in epigenetics and miRNA pattern lead to progression from precancerous lesions to HCC [29,30]. The epigenetic alterations including DNA hypermethylation or hypomethylation, dysregulation of histone modification patterns, chromatin remodeling *etc* are associated with HCC [31]. Tumor suppressor genes such as *SOCS1, hMLH1, GSTP1, MGMT, CDH1* and *TIMP3 etc* are inactivated by the virtue of promoter hypermethylation [32]. Detection of cancer at an early stage is vital to reduce the mortality rate by providing appropriate treatment based on cancer stage. In previous studies, researchers focused mainly on identification of differentially expressed RNA transcripts or genes in LIHC. Best of our knowledge, no method has been developed for predicting stage of LIHC from genomic profile of samples. In this study, a systematic attempt has been made to develop model for discriminating early and late stage of LIHC samples. Firstly, we identified CpG sites that are differentially methylated in early and late stage of LIHC. Secondly, we identified aberrantly expressed RNA transcripts that can differentiate early stage from late stage of LIHC. Thirdly, models were developed based on different machine learning techniques for predicting stage of LIHC samples using the above derived genomic and epigenomic features. Using diverse feature spaces, we were able to establish models discriminating early and late stage of LIHC cancer. In addition, we made an attempt to develop models for discriminating normal and LIHC tissue samples. Our models successfully predict LIHC samples with high accuracy. It indicates that it is easy to predict LIHC samples but it is challenging to classify them in early and late stage. We also attempted to develop multiclass prediction models to classify samples in three classes; i) normal or control samples, ii) LIHC early stage and iii) LIHC late stage tissue samples.

## Data Description

### Main Dataset of Early & Late Stage Samples

We extracted the expression and methylation profiles of Liver Hepatocellular Carcinoma (LIHC) samples from GDC Data Portal (https://portal.gdc.cancer.gov/). In addition, manifest, metadata, clinical data, biospecimen files were downloaded to obtain clinical information using Biospecimen Core Resource (BCR) IDs of patients. Finally, we obtained 173 stage-I, 87 stage-II, 85 stage-III and 5 stage-IV stage samples. Clinical characteristics of these patients displayed in figure S1 (Additional file 3, Figure S1). As the number of stage-IV samples in the dataset is small, we have considered stage-II, stage-III and stage-IV samples as late stage samples, while stage-I samples as early stage samples as stage I samples are of localized cancer which shows no sign of metastasis. We also downloaded the methylation profiles acquired using the Illumina Human-Methylation450K DNA Analysis BeadChip assay, based on genotyping of bisulfite-converted genomic DNA at individual CpG-sites. This provides Beta values, a quantitative measure of DNA methylation [33]. In addition, for each subject, RNA expression in terms of FPKM values for 60,483 RNA transcripts were reported. In this study, we have used FPKM values of RNA transcripts as quantification values.

## Analyses

### Models for classification of early staee and late staee of LIHC samples

Our primary objective is to identify potential markers *i.e.* CpG sites and RNA transcripts that can classify early stage tissue samples from late stage tissue samples. Subsequently in silico predictive models were developed based on these signature markers using various machine learning algorithms. Potential markers and prediction models based on them are explained in the following sections. The overall work flow represented in Figure 1.

**Figure 1:**
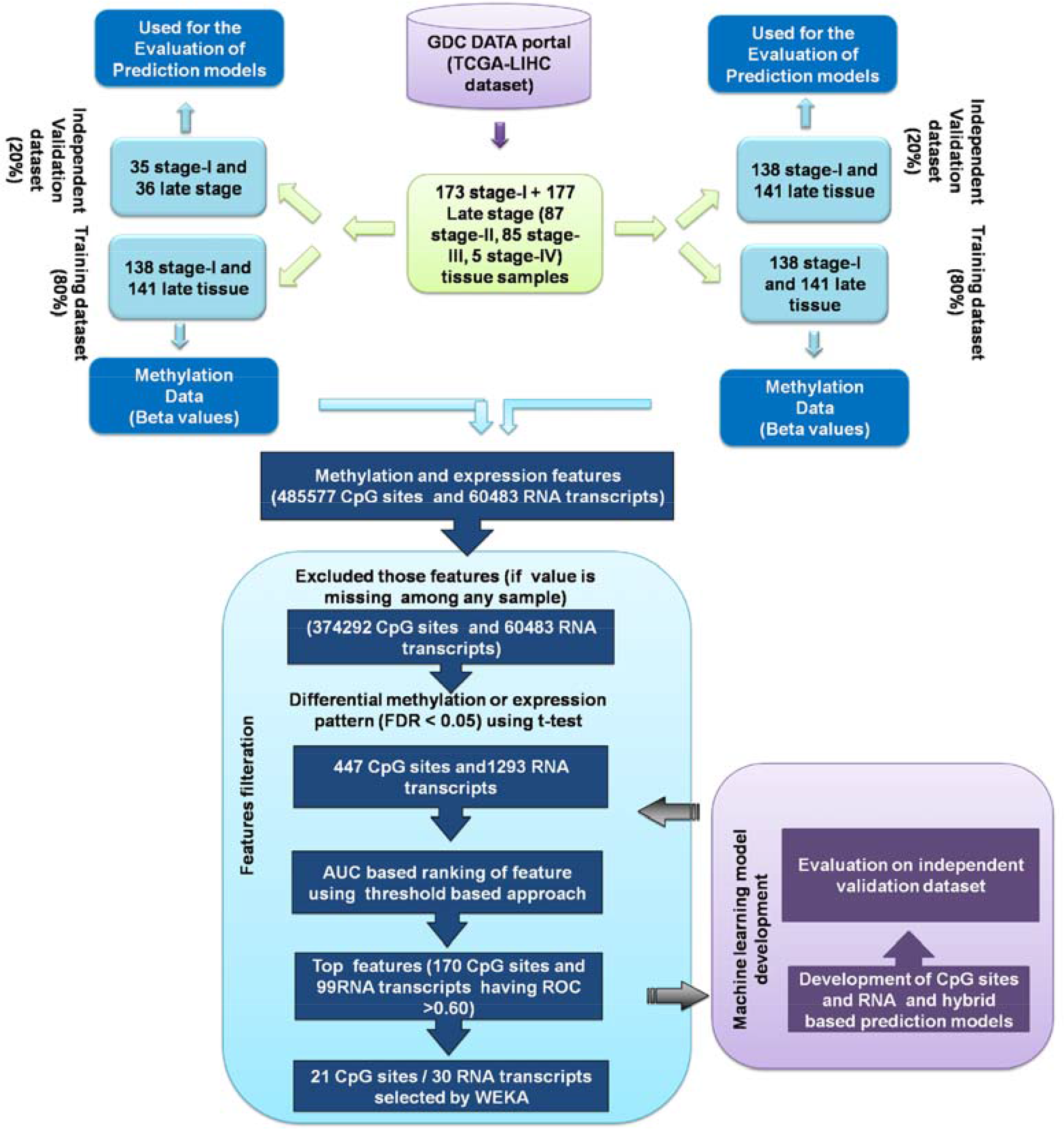
The workflow representing the analysis of methylation and expression profile of LIHC early and late stage samples

#### Single feature based stage classification model using CpG sites

Here, we used single feature based classification technique to classify early and late stage samples. In this study, we first identified 447 CpG sites (from 3,74,292 sites) that are significantly differentially methylated in early and late stage of sample using t-test (FDR<0.05). These differentially methylated sites are further segregated in two classes; i) 199 hypermethylated CpG sites (high level of methylation in early stage) and ii) 248 hypomethylated CpG site (low level methylation in early stage). Further, we used each differentially methylated CpG sites to develop a simple threshold-based model for classifying early and late stage samples. In the threshold-based model, a sample is classified in the early stage if the methylation level of CpG site (in case site is hypermethylated) is higher than a threshold value otherwise in late stage. In these models, the threshold is varied incrementally from minimum to maximum beta-value. In the final step, that threshold is selected which leads to maximum AUROC between early and late samples. Subsequently, all the 447 CpG sites are ranked according to AUROC to assess the ability of a CpG site to categorize early and late stage tissue samples (Additional file 1, Table_S1). As shown in Table_S1 (Additional file 1), there are 170 differentially methylated CpG sites (LS-CPG-AUROC) that can differentiate two types of samples with high precision (ROC ≥ 0.6). LS-CPG-AUROC further includes 105 hypermethylated and 65 hypomethylated CpG sites. The differential methylation pattern can be observed in heatmap (Additional file 4, Figure_S2). Hypermethylated CpG sites in early stage such as cg12595697, cg11232136, cg01055099 can distinguish early and late stage samples with AUROC 0.66, 0.65 and 0.65 at threshold 0.94, 0.90 and 0.84 respectively. It indicates that if the beta value of these CpG sites is greater than the mentioned threshold, then sample belongs to early stage, otherwise to the late stage. The hypomethylated sites in the early stage such as cg07132710 and cg22491320 can classify early stage samples from late stage with AUROC at 0.65 at threshold 0.63 and 0.43. It signifies that if the beta value of these CpG sites is lower than the mentioned threshold, then the sample belongs to early stage, otherwise late stage. Methylation pattern of the top 20 CpG from 447 CpG sites is shown **Figure 2A** and their chromosome location and associated genes are represented in **Figure 2B**.

**Figure 2:**
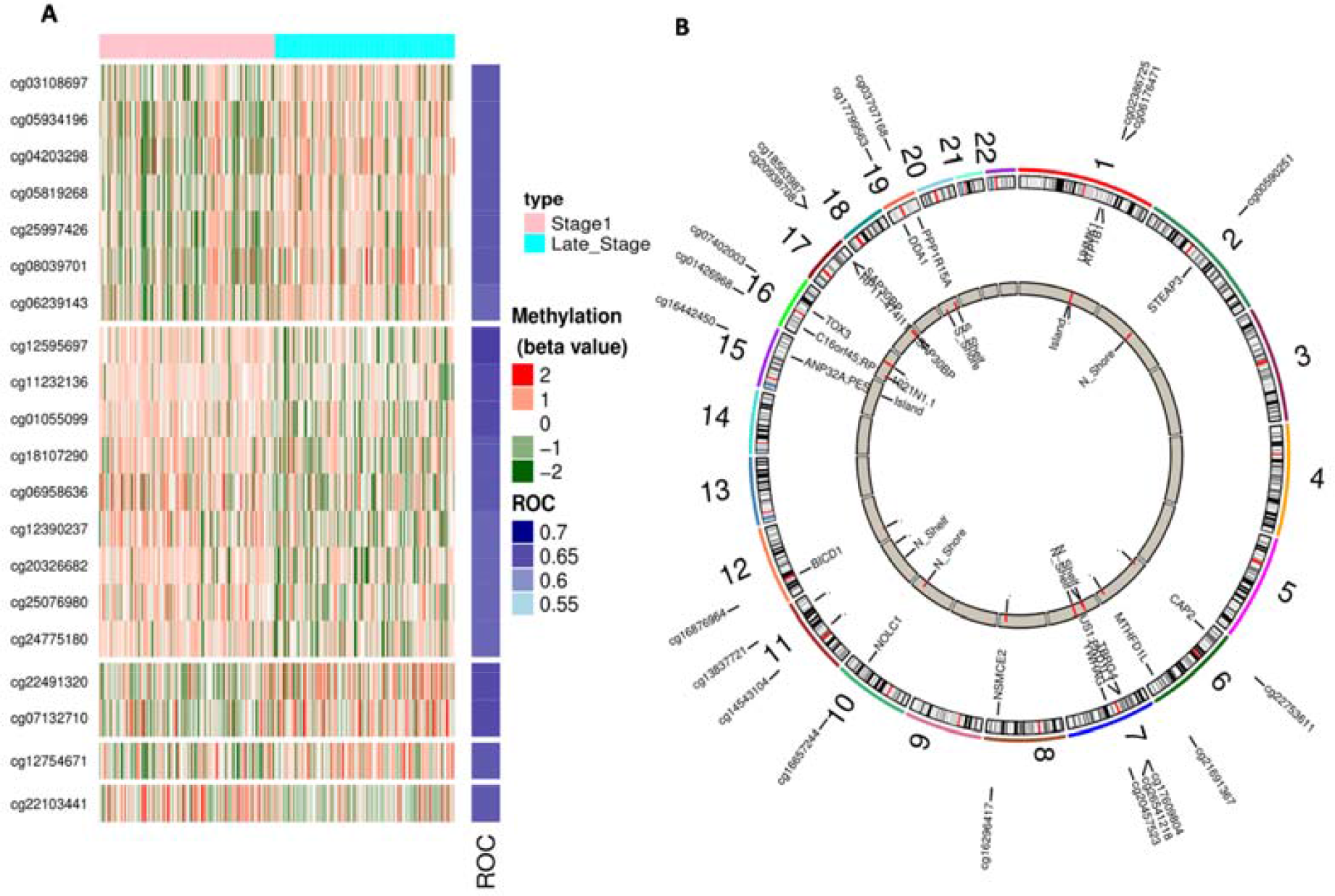
A) Heatmap displaying the differential methylation pattern (with FDR < 0.05) and B) Circos plot representing the chromosome location of top 20 CpG sites (AUROC based) in early versus late stage of LIHC.

LS-CPG-AUROC signature is significantly enriched (adjusted *p*-value <0.05) in various immune system associated pathways and cancer associated cell signaling pathways present in BioCarta. For instance, 4 out of 55 genes of T Cell Receptor Signaling Pathway and 5 out of 33 genes of Integrin Signaling Pathway_Homosapiens, are enriched in LS-CPG-AUROC. Furthermore, *ITGB1, SHC1* and *PTK2* are associated with PTEN dependent cell cycle arrest and apoptosis, VEGF, Hypoxia, and AngiogenesisPathway. This signature is also enriched in actin filament binding (GO:0051015).

#### Multiple features based stage classification model using CpG sites

One of the challenges in classification models is to improve the performance of models. Though, we got a single CpG site like cg12595697, which can classify samples with AUROC 0.66. To utilize information from multiple features, we developed models using machine learning techniques. Firstly, models developed based on various machine learning techniques using above LS-CPG-AUROC. As shown in Table_S2 (Additional file 1), most of the models attain reasonable high performance on training dataset, maximum AUROC 0.79 for SVM, but poor AUROC *i.e.* 0.66 on external validation dataset.

Therefore, we selected 21 features using WEKA from 447 CpG differentially methylated sites with FDR (False discovery Rate) < 0.05 for early stage v/s late stage. These 21 selected features (CpG sites) named as LS-CPG-WEKA were used to develop models for classification using different machine learning techniques (Table 1). The SVM based model achieved the highest accuracy of 75.99% with AUROC 0.81andaccuracy of 73.24% with AUROC 0.78 on the training dataset and the validation dataset respectively as shown in Table 1. Models based on Random forest, SMO and Naïve Bayes also achieved performance in the similar range. Among 21 CpG sites (selected features), 9 and 12 were hypomethylated and hypermethylated CpG sites respectively. Heatmap displaying the methylation pattern (Figure 3A) and Circos plot representing the chromosome location, genes associated with them and feature type shown in Figure 3B.

**Table 1:** The performance of different machine learning techniques based stage classification models developed using 21 methylation CpG sites selected by WEKA (LS-CPG-WEKA).

| Machine Learning Techniques | Dataset | Performance Measures |  |  |  |  |
| --- | --- | --- | --- | --- | --- | --- |
|  |  | Sensitivity | Specificity | Accuracy (%) | MCC | AUROC |
| SVM | Training | 80.43 | 71.63 | 75.99 | 0.52 | 0.81 |
|  | Validation | 71.43 | 75 | 73.24 | 0.46 | 0.78 |
| Random Forest | Training | 77.54 | 73.05 | 75.27 | 0.51 | 0.8 |
|  | Validation | 71.43 | 83.33 | 77.46 | 0.55 | 0.79 |
| Naïve Bayes | Training | 87.68 | 62.41 | 74.91 | 0.52 | 0.82 |
|  | Validation | 80 | 63.89 | 71.83 | 0.44 | 0.82 |
| SMO | Training | 82.61 | 69.5 | 75.99 | 0.53 | 0.76 |
|  | Validation | 74.29 | 75 | 74.65 | 0.49 | 0.75 |
| J48 | Training | 62.32 | 75.18 | 68.82 | 0.38 | 0.67 |
|  | Validation | 45.71 | 61.11 | 53.52 | 0.07 | 0.57 |

**Figure 3:**
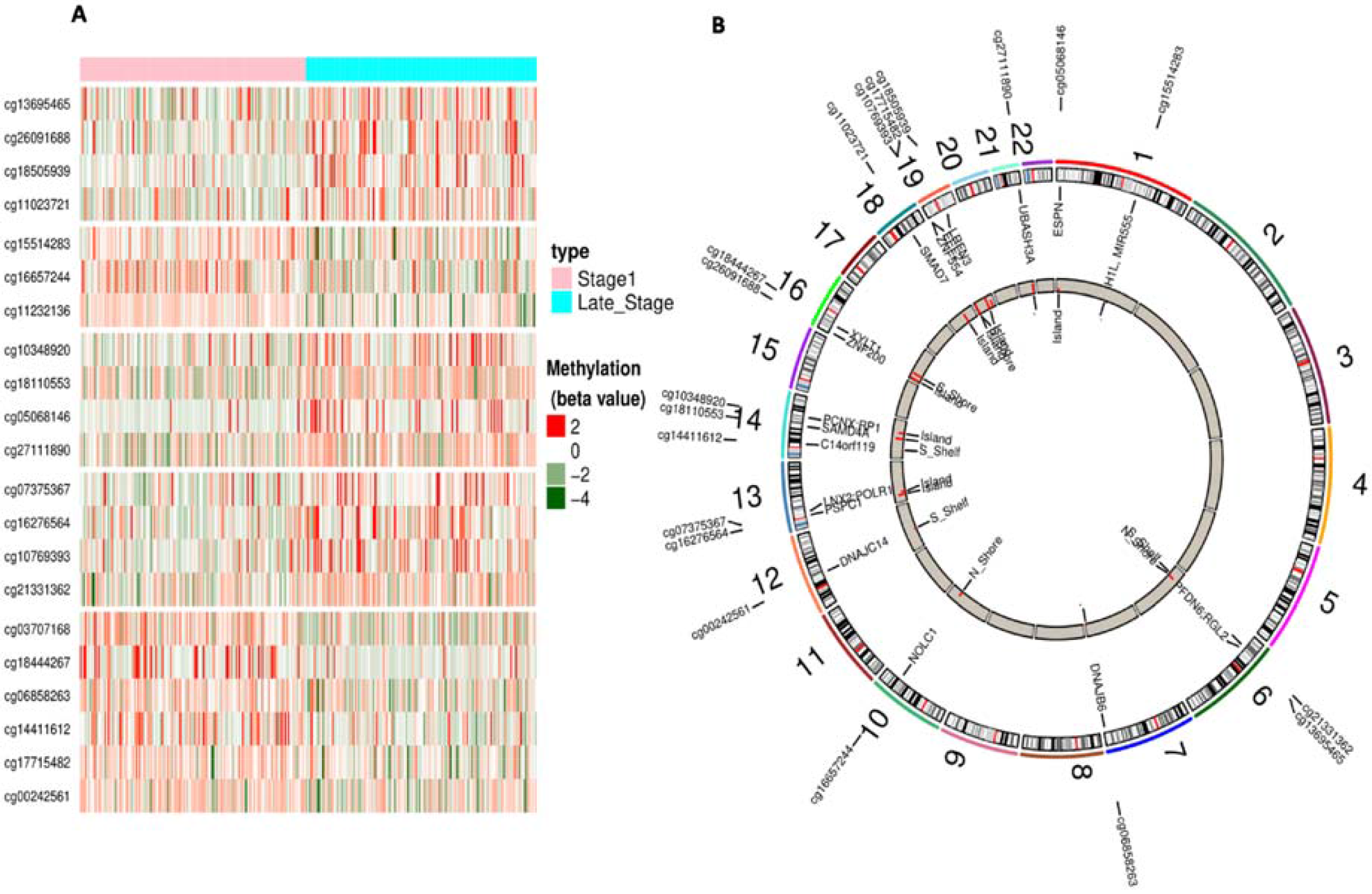
A) Heatmap displaying the differential methylation pattern (with FDR < 0.05) and B) Circos plot representing the chromosome location of top 21 CpG sites (LS-CPG-WEKA) in early versus late stage of LIHC

Interestingly 9 hypermethylated CpG sites of LS-CPG-WEKA signature in early stage are associated with 12 genes (*EEF2, NOLC1, PPP1R15A, DNAJC14, DNAJB6, XYLT1* etc.) that are significantly enriched (adjusted p-value <0.05) in various GO biological processes such as two genes are involved in positive regulation of translation (GO:0045727) and rescue of stalled ribosome (GO:0072344). While, hypomethylated CpG sites are associated with 14 genes (*SMAD7, PCNX, RGL2, POLR1D, ZNF554, ZNF200 etc*) and these genes are significantly enriched in different BioCarta pathways. For instance, *SMAD7*, a component of catenin complex (GO:0016342), is involved in TGF beta signaling pathway_Homosapiens_h_tgfbPathway and has been shown to inhibit TGF-beta (Transforming growth factor) and active in signaling by associating with their receptors. *ZNF554, PLOR1D* and *PSPC1* are involved in regulation of transcriptional activity.

#### Stage classification models using the expression of RNA transcripts

In above section, we studied methylation level of CpG sites in early and late stage samples as well as the classification models developed using methylation of CpG sites. Similarly, in this section, we analyzed expression of RNA transcripts in both types of samples as well as developed stage classification models. Firstly, RNA transcripts number was reduced to 103 (LS-RNA-AUROC) after applying stringent criteria (FDR adjusted p-value < 0.01) from 60,483. In LS-RNA-AUROC signature, 39 transcripts were over-expressed and 64 transcripts were under expressed in early stage LIHC samples. Secondly, we developed single feature based models using each transcript, similar to single features based models for CpG sites. Finally, we ranked RNA transcripts based on AUROC to discriminate early stage from late stage tissue samples. The 99 RNA transcript having AUROC greater or equal to 0.6 is shown in Table_S3 (Additional file 1, Table S3) with their expression patterns illustrated in Figure_S3 (Additional file 5). Furthermore, *NCAPH, CYP4A22, HSD17B6, GLYATL1* and *FTCD* are top 5 RNA transcripts which distinguish early stage and late stage LIHC samples with AUROC >= 0.65. Among them, *NCAPH* is top performer with AUROC 0.66 at threshold 1.45. It means that if the log2-FPKM value of *NCAPHis* lower than 1.45, then the sample belongs to early stage otherwise to late stage as it is under-expressed in early stage. The expression pattern of top 20 genes in early stage samples in comparison to late stage tissue samples displayed in Figure 4A.

**Figure 4:**
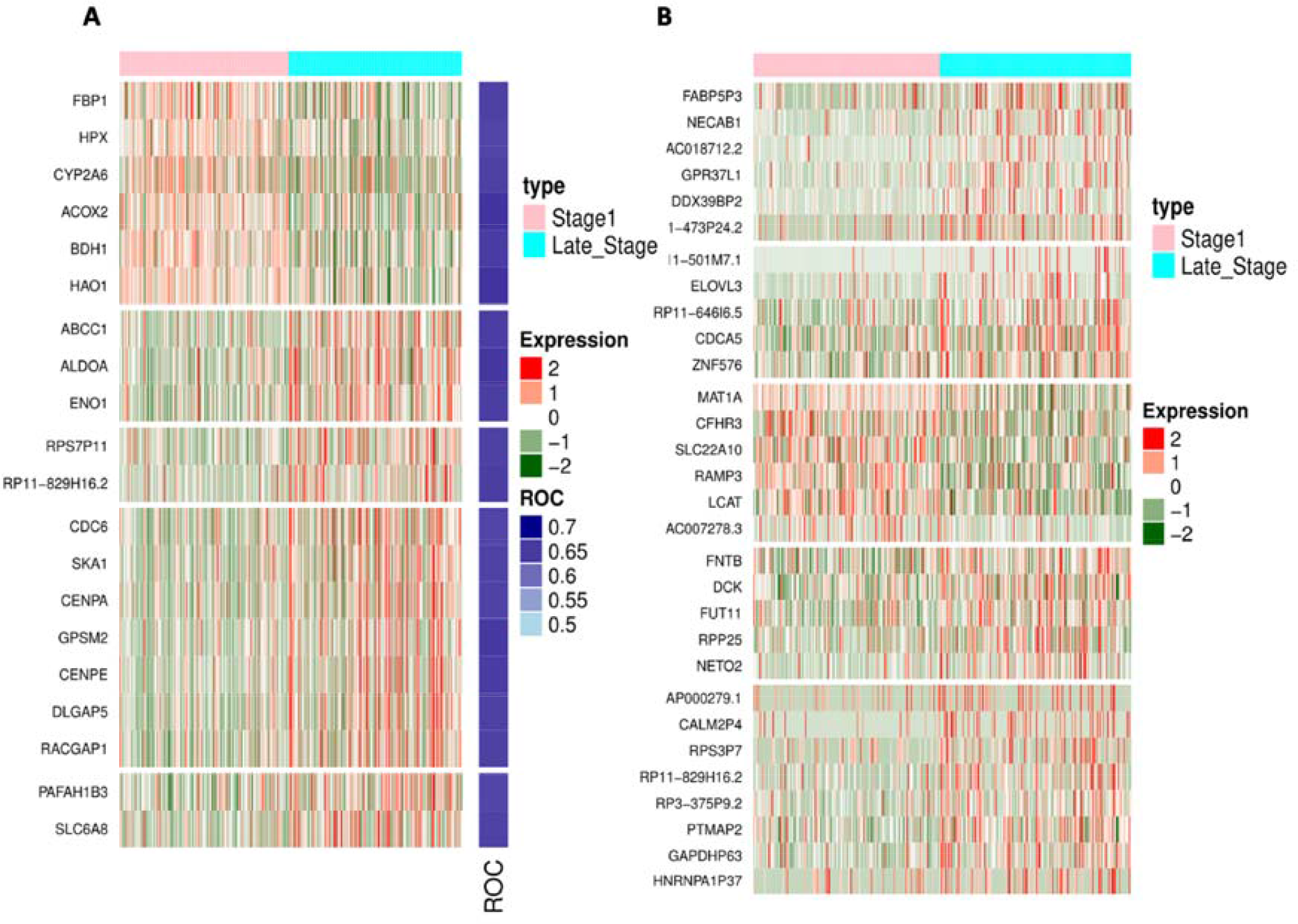
The differential expression pattern of A) top 20 RNA transcripts (AUROC based) and B) 30 RNA transcripts (LS-RNA-WEKA) in early stage versus late stage tissue samples (With FDR <0.01).

Gene enrichment analysis of LS-RNA-AUROC which includes 61 downregulated and 39 upregulated transcripts in early stage of LIHC using Enrichr displayed in Figure 5 (A and B). The enrichment analysis of these transcripts indicates that downregulated transcripts are mostly enriched in metabolic processes related pathways; whereas, upregulated transcripts are associated with pathways that involved in normal functioning of liver. Further, downregulated transcripts enriched in cell cycle GO terms that contribute to cell growth while upregulated transcripts enriched in GO terms associated with common functioning of liver. In addition downregulated transcripts enriched in MSigDB oncogenic signatures i.e. RPS14_DN.V1_DN and CSR_LATE_UP.V1_UP, while upregulated transcripts enriched in PKCA_DN.V1_UP.

**Figure 5:**
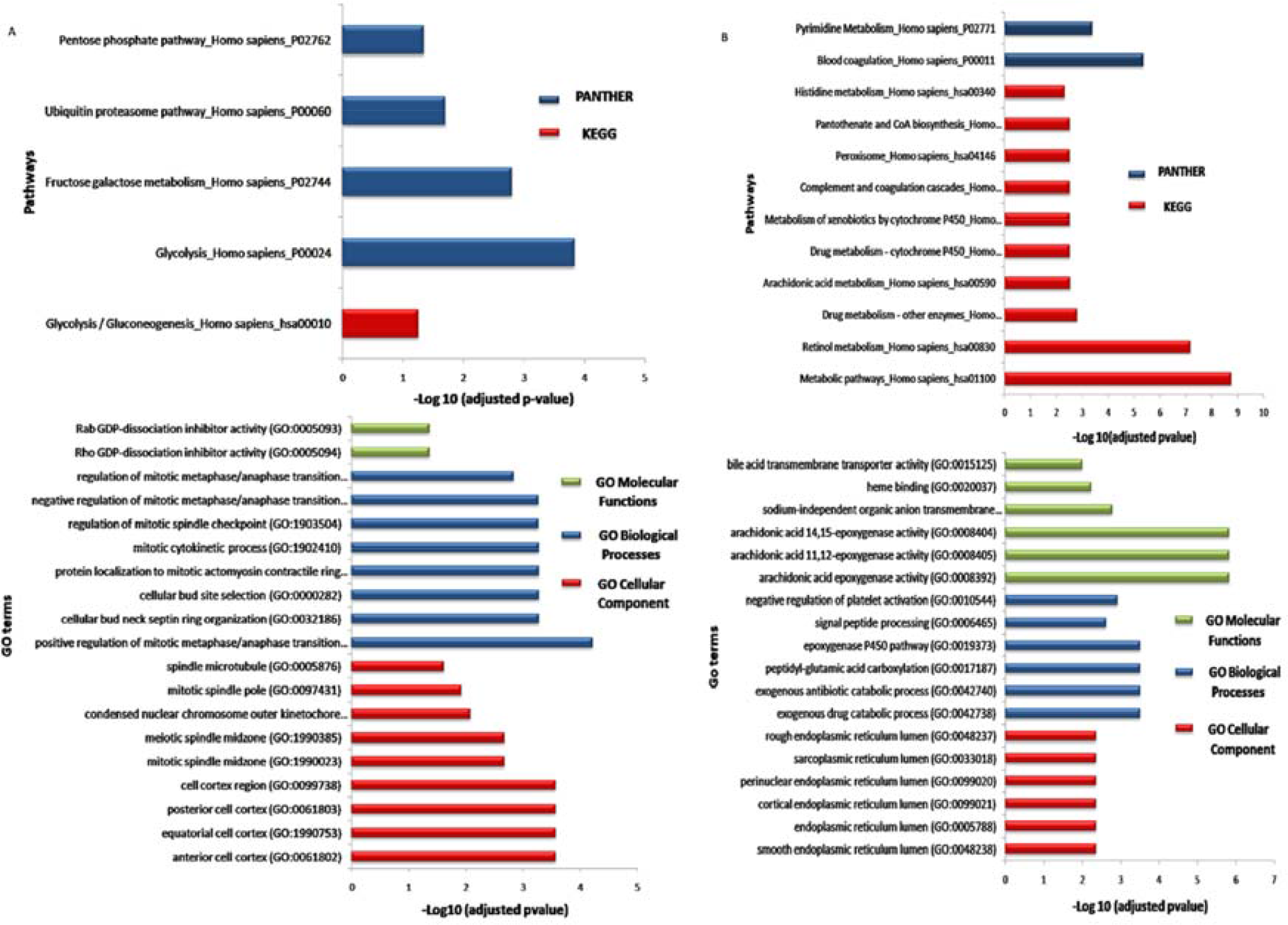
Gene Enrichment analysis of LS-RNA-AUROC, A) 61 downregulated and B) 39 upregulated RNA transcripts in early stage in comparison to late stage of LIHC.

#### Multiple features based stage classification model using RNA transcripts

In addition to single feature based models, we have also developed models using expression of multiple RNA transcripts implementing different machine learning techniques. First, 30 RNA transcripts (named as LS-RNA-WEKA) selected from 1,293 RNA transcript using WEKA, subsequently prediction models developed for stage classification. As shown in Table 2, the Naive Bayes based model achieved maximum accuracy of 74.55% with AUROC 0.79 on training dataset and accuracy of 76.06% with AUROC 0.79 on validation dataset respectively (Table 2). Among 30-signature RNA transcripts (LS-RNA-WEKA), 24 RNA transcripts have been found to be underexpressed, while 6 observed as overexpressed in early stage (Additional file 1, Table S4). Further among 30 transcripts,15 are protein coding genes which include *CSNK1D*, *DCK*, *ZNF576*, *RPP25*, *SLC22A10*, *FNTB*, *LCAT*, *MAT1A* and *CDCA5 etc,* 9 processed pseudogenesinclude GAPDHP63, RP11-829H16.2, AC018712.2 *etc*, 2 unprocessed pseudogenes, 1 each of lincRNA, miRNA,sense_intronic transcript. Figure 4B represents the expression pattern of these RNA transcripts among early and late stage tissue samples. In addition, we also developed prediction model using 100 features selected by F-ANOVA method. The performance (accuracy 70.42, AUROC 0.74 on independent validation dataset) of model has decreased with increasing the number of features as shown in Table_S5 (Additional file 1).

It has been observed that most of overexpressed genes such as *CDCA5*, *CSNK1D*, *DCK*, *ZNF576* and *RPP25 etc* in late stage are involved in cell growth promotion processes. While underexpressed genes such as *LCAT*, *CFHR3*, *RAMP3*, *MAT1A etc* associated with normal functioning of liver and immune functions as shown in Table_S6 (Additional file 1).

**Table 2:**
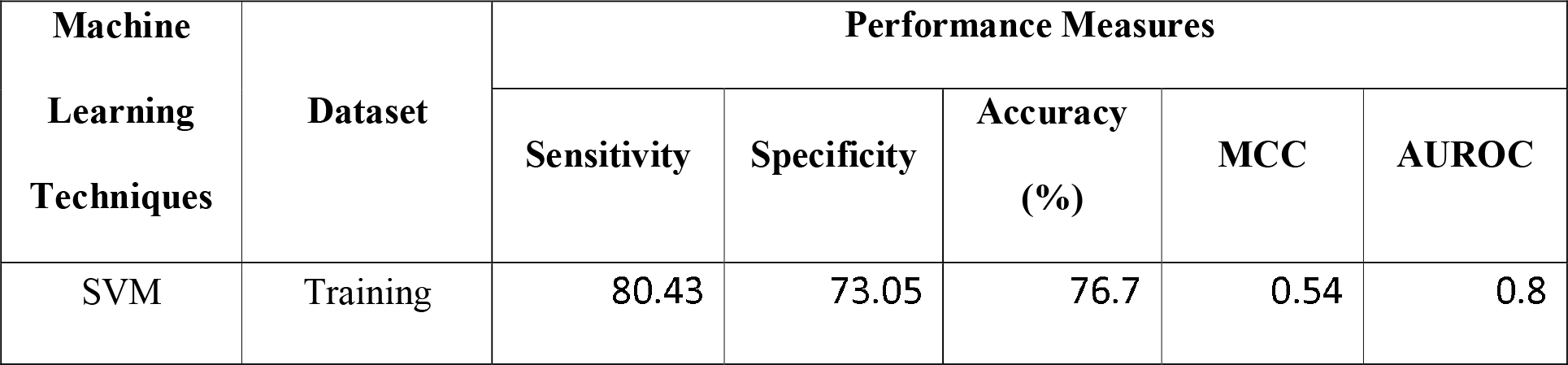
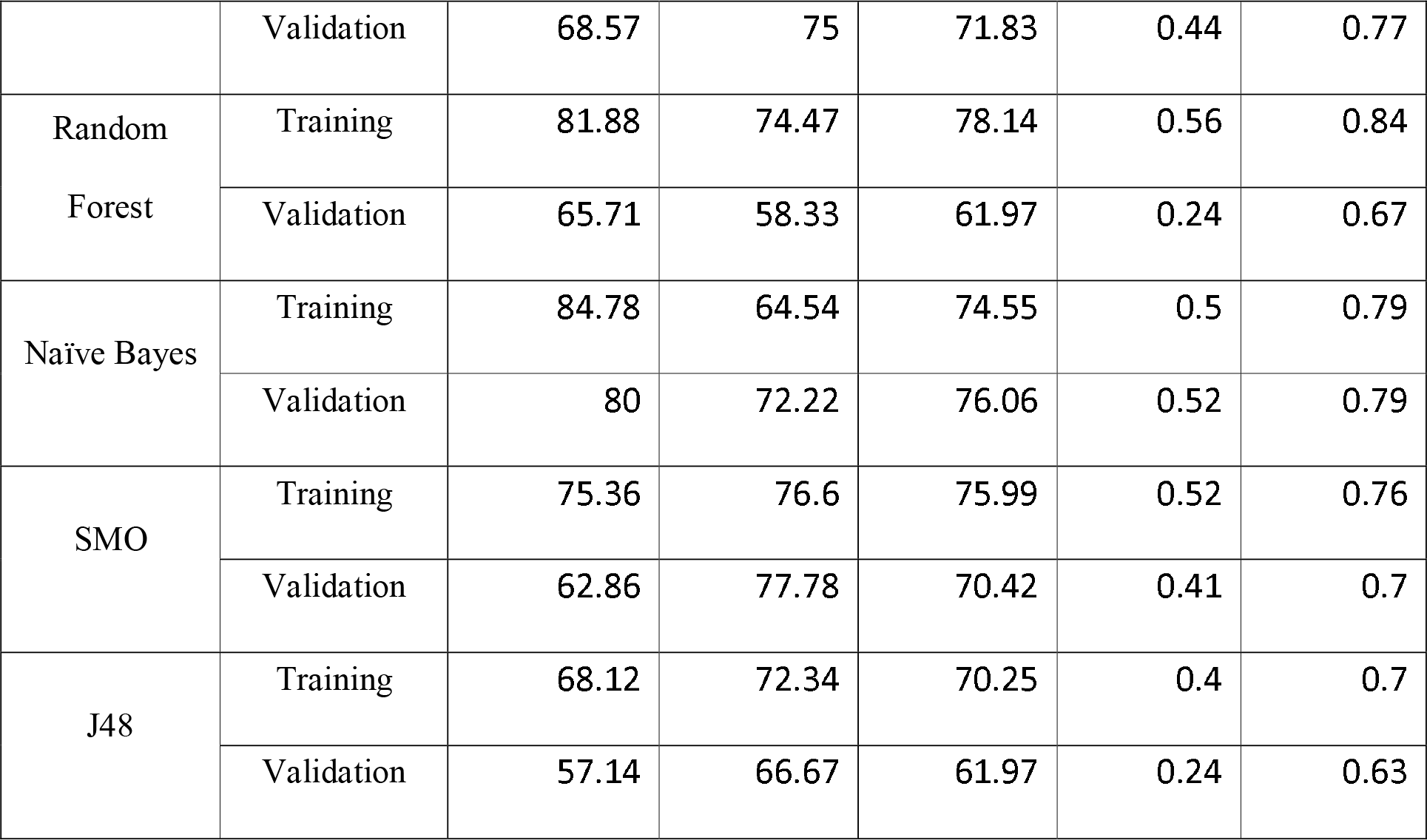
The performance of stage classification models developed using 30 RNA transcripts selected using WEKA from 103 RNA transcripts (LS-RNA-WEKA).

#### Stage classification models using hybrid features

In addition to single type of features (CpG methylation or RNA transcripts), we also developed model by combining both type of features. Prediction models were developed using 51 features (21 CpG sites and 30 RNA transcripts) selected by WEKA named as LS-CpG-RNA-hybrid to classify early and late stage tissue samples. The model developed using Naïve Bayes based model achieved the highest accuracy 78.14% with AUROC 0.81 on the training dataset and accuracy 78.87% and 0.82 on the independent dataset (Table 3). Also, we also developed models using selected 38 features (15CpG sites and 23RNA transcripts) obtained from 1740 features (1293 RNA transcripts and 447CpG sites), (Additional file 1, Table_S7). In summary, we got best performance of Naïve Bayes based model developed using 51 hybrid features (21 CpG sites and 30 RNA transcripts). In spite of the best efforts, we were capable to discriminate the early and late stage samples with maximum AUROC of 0.82.

**Table 3:**
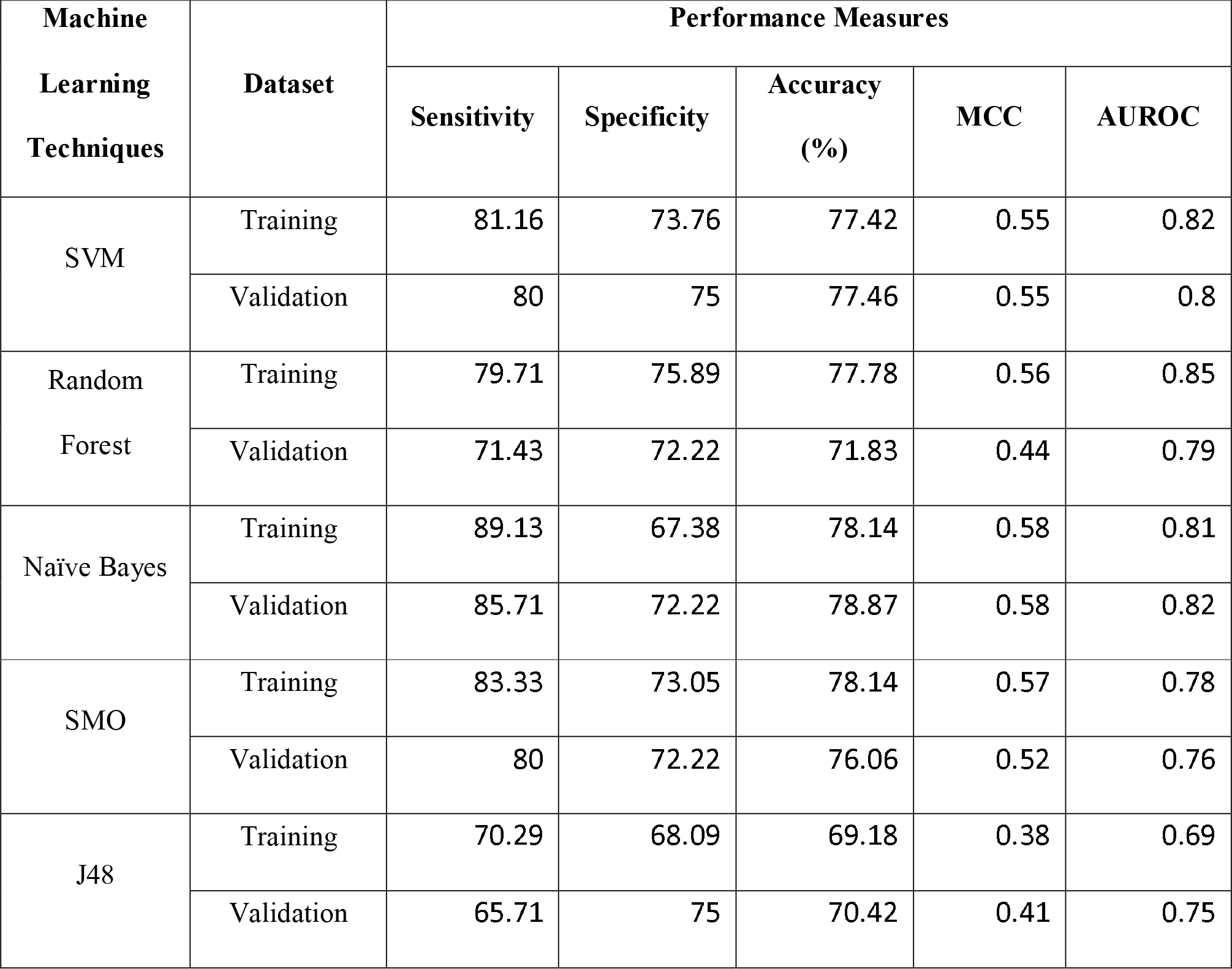
The performance of stage classification hybrid models developed using 51 features (LS-CpG-RNA-hybrid) that comprise 21 CpG sites and 30 RNA transcripts.

| Machine Learning Techniques | Dataset | Performance Measures |  |  |  |  |
| --- | --- | --- | --- | --- | --- | --- |
|  |  | Sensitivity | Specificity | Accuracy (%) | MCC | AUROC |
| SVM | Training | 81.16 | 73.76 | 77.42 | 0.55 | 0.82 |
|  | Validation | 80 | 75 | 77.46 | 0.55 | 0.8 |
| Random Forest | Training | 79.71 | 75.89 | 77.78 | 0.56 | 0.85 |
|  | Validation | 71.43 | 72.22 | 71.83 | 0.44 | 0.79 |
| Naïve Bayes | Training | 89.13 | 67.38 | 78.14 | 0.58 | 0.81 |
|  | Validation | 85.71 | 72.22 | 78.87 | 0.58 | 0.82 |
| SMO | Training | 83.33 | 73.05 | 78.14 | 0.57 | 0.78 |
|  | Validation | 80 | 72.22 | 76.06 | 0.52 | 0.76 |
| J48 | Training | 70.29 | 68.09 | 69.18 | 0.38 | 0.69 |
|  | Validation | 65.71 | 75 | 70.42 | 0.41 | 0.75 |

#### Models for classification of LIHC and normal samples

##### Models for classification of LIHC and normal samples using CpG sites

In this study, we also developed models for discriminating LIHC and normal tissue samples using genomic and epigenetic profiles. These models were developed on 424 samples (50 normal and 374 LIHC), where training dataset consists of 339 samples (40 normal and 299 LIHC) and independent dataset consists of 85 samples (10 normal and 75 LIHC). We identified 1,386 CpG sites that are differentially methylated in LIHC and normal tissue samples (minimum mean difference of methylation beta values is 0.4) with Bonferroni *p*-value < 0.0001. These 1,386 sites contain 125 hypermethylated and 1,261 hypomethylated in cancer samples (Additional file 1, Table_S8). Further, we developed single feature based models using these CpG sites and ranked them based on performance of models. Top 496 CpG sites named as LCN-CpG-AUROC (68 hypemethylated and 428 hypomethylatedCpG sites in LIHC) with AUROC equal or greater than 0.9 enlisted in (Additional file 1, Table_S9). The methylation pattern of top 496CpG sites in cancer and normal tissue samples is shown in Figure_S4 (Additional file 6). We have obtained 14 CpG sites that can discriminate LIHC and normal samples with precision (ROC >= 0.95). Single feature based models developed using CpG site cg07274716, cg24035245 and cg20172627 were able to classify normal and LIHC tissue with AUROC 0.97, 0.96 and 0.95 at threshold 0.35, 0.44, 0.32 and 0.12 respectively. Methylation pattern of top 10 CpG sites is shown by Heatmap (Additional file 7, Figure_S5A) and chromosome locations and associated genes of the CpG sites are represented in Circos plot (Additional file 8, Figure_S6).

In addition to the single feature based models, we also developed models using multiple features to classify LIHC and normal samples. Our SVM based model developed using 104 CpG sites (selected using WEKA from 496CpG sites) achieves maximum accuracy of 98-99 % and AUROC 0.99 on independent and training dataset. We further reduce features and developed models using 100, 50, 25, 20, 10 and 5 features or CpG sites (selected using F-ANOVA), (data not shown). Our models based on even small number of features (*i.e.* 5 CpG sites named as LCN-5-CpG) got reasonably high performance (ROC ~0.99). Models based on Random Forest, Naive Bayes, SMO algorithms performed reasonably good with accuracy 95%-97%, and AUROC of 0.94 - 0.97 on training dataset and accuracy 96 - 97% and AUROC of 0.94 - 99 on an independent validation dataset (Table 4). This observation shows that LIHC samples can be discriminated with high precision even using small number of CpG methylation based biomarkers.

**Table 4:** The performance models developed for discriminating LIHC and normal
samples, using 5 methylation CpG sites.

| Machine Learning Techniques | Dataset | Performance Measures |  |  |  |  |
| --- | --- | --- | --- | --- | --- | --- |
|  |  | Sensitivity | Specificity | Accuracy (%) | MCC | AUROC |
| SVM | Training | 99.33 | 90 | 98.23 | 0.91 | 0.99 |
|  | Validation | 98.67 | 90 | 97.65 | 0.89 | 0.99 |
| Random Forest | Training | 99 | 87.5 | 97.64 | 0.88 | 0.97 |
|  | Validation | 98.67 | 90 | 97.65 | 0.89 | 0.99 |
| Naïve Bayes | Training | 96.66 | 90 | 95.87 | 0.82 | 0.94 |
|  | Validation | 97.33 | 90 | 96.47 | 0.84 | 0.95 |
| SMO | Training | 99 | 90 | 97.94 | 0.9 | 0.94 |
|  | Validation | 98.67 | 90 | 97.65 | 0.89 | 0.94 |
| J48 | Training | 97.66 | 80 | 95.58 | 0.79 | 0.86 |
|  | Validation | 98.67 | 80 | 96.47 | 0.82 | 0.89 |

To detect the biological relevance of the genes associated with these signature CpG sites (LCN-5-CpG), enrichment analysis performed using Enrichr. This analysis indicates *MYH9* is significantly involved in molecular functions such as actin-dependent ATPase activity (GO:0030898), microfilament motor activity (GO:0060002, GO:0060001). It is also involved in GO biological processes such as positive regulation of protein processing in phagocytic vesicle (GO:1903923), PMA-inducible membrane protein ectodomain proteolysis (GO:0051088), malonyl-CoA metabolic process (GO:2001293). *ACSF2* is involved in acyl-CoA metabolic process (GO:0006637), malonyl-CoA metabolic process (GO:2001293), *etc.* Furthermore, the enrichment analysis signifies the involvement of *MYH9* in Nicotinic acetylcholine receptor signaling pathway, Cytoskeletal regulation by Rho GTPase, Inflammation mediated by chemokine and cytokine signaling pathway_Homo sapiens_P00031 PANTHER pathways. In addition, Enrichr suggest the enrichment of *ACSF2* in LEF1 _UP.V1 _DN, while *CHAD* involved in AKT_UP_MTOR_DN.V1 _DN, and AKT_UP.V1_DN MSigDB Oncogenic Signatures set (*p*-value <0.05)

##### Models for classification of LIHC and normal samples using RNA transcripts

Moreover, we have also identified 147 RNA transcripts that are significantly differentially expressed in LIHC and normal samples (Additional file 1, Table_S10). The performance of models developed using top 53 RNA transcripts (ROC equal or greater than 0.9) is shown in Table_S11 (Additional file 1). The expression pattern of top 53RNA transcripts (named as LCN-RNA-AUROC) in cancer and normal tissue samples have been shown in heatmap (Additional file 9, Figure_S7). Among 53 transcripts, 50 are protein coding, one each of Linc-RNA, snoRNA and processed transcripts. Further, among them 18 and 35 observed to be overexpressed and underexpressed respectively in LIHC in comparison to normal tissue samples. Moreover, *CLEC4G*, *CLEC4M*, *FCN2*, *COLECIO*, *CLEC1B* and *PLVAP* can classify LIHC and normal tissue samples with AUROC >= 0.96. *CLEC4G* which is underexpressed gene in cancer is chief performer with AUROC ~0.99 at threshold 2.9. It means if the log2 of FPKM value of *CLEC4G* is less than 2.9 then sample belongs to cancer otherwise normal. *PLVAP* is overexpressed gene in cancer samples, it can distinguish cancer and normal samples with AUROC 0.97 at threshold 3.8. It suggested if the log2 of FPKM value of *PLVAP* above 3.8 then the sample is cancer otherwise normal. Expression pattern of top 10 RNA transcripts displayed in heatmap (Additional file 7, Figure_S5B)

Gene enrichment analysis of LCN-RNA-AUROC is represented in Figure_S8A and Figure_S8B (Additional file 10). This functional enrichment of these genes suggested their involvement in various growth progression events or cancer development processes. As these genes upregulated in LIHC or cancerous condition; this might lay insight toward their contribution in the diseased state. While the underexpressed genes in LIHC are enriched in processes and pathways associated with normal activity or functioning of liver. Moreover they are also enriched in various MSigDB oncogenic signatures (Additional file 1, Table_S12).

Further, we also proceed to develop multiple genes based models that can outperform single gene based model. Thus we developed models based on multiple RNA transcripts/features selected by different techniques i.e., WEKA and F-ANOVA method. In this, an attempt has been made to develop prediction model using 35 WEKA selected features (from 147 RNA transcripts) to distinguish liver cancer samples from the normal samples. The SVM based model attains maximum accuracy nearly 99% with AUROC 0.99on both training and independent validation dataset respectively (result not shown). To develop the prediction model based on least number of genes/features, that can categorize LIHC samples from normal tissue samples with reasonably good accuracy; we firstly selected 25, 20, 15, 10 and 5 feature sets (results not shown) using F-ANOVA method. Then we developed prediction models on training dataset and independent validation dataset. The model’s performance on independent validation dataset is similar to the performance obtained using CpG sites. Interestingly models based on these feature sets almost performed comparably to that of model based on 35 features selected by WEKA. 5-RNA feature set (LCN-5-RNA) based SVM prediction model categorize samples with accuracy 98.53 and AUROC 0.97 of training dataset and with accuracy 97.65 and AUROC 0.93 of the independent validation dataset. In addition models based on the Random forest, SMO, Naïve Bayes and J48 algorithms also performed almost equally to that of SVM as indicated in Table 5.

**Table 5:** The performance of models developed for discriminating LIHC and normal samples using 5 RNA transcripts.

| Machine Learning Techniques | Dataset | Performance Measures |  |  |  |  |
| --- | --- | --- | --- | --- | --- | --- |
|  |  | Sensitivity | Specificity | Accuracy (%) | MCC | AUROC |
| SVM | Training | 97.32 | 97.5 | 97.35 | 0.89 | 0.99 |
|  | Validation | 97.33 | 90 | 96.47 | 0.84 | 0.97 |
| Random Forest | Training | 97.66 | 90 | 96.76 | 0.85 | 0.98 |
|  | Validation | 98.67 | 80 | 96.47 | 0.82 | 0.89 |
| Naïve Bayes | Training | 96.66 | 97.5 | 96.76 | 0.86 | 0.97 |
|  | Validation | 97.33 | 90 | 96.47 | 0.84 | 0.94 |
| SMO | Training | 97.32 | 97.5 | 97.35 | 0.89 | 0.97 |
|  | Validation | 97.33 | 90 | 96.47 | 0.84 | 0.94 |
| J48 | Training | 96.99 | 97.5 | 97.05 | 0.87 | 0.96 |
|  | Validation | 98.67 | 90 | 97.65 | 0.89 | 0.94 |

Enrichment analysis of LCN-5-RNA using Enrichr implies the involvement of *CLEC4M* in Phagosome KEGG pathways, and *GDF2* in the TGF-beta signaling pathway_Homo sapiens_P00052 (p-value <0.05). *CLEC4M* involved in receptor-mediated virion attachment to host cell (GO:0046813), evasion or tolerance of host defenses by virus (GO:0019049) *etc.* This analysis also denotes their enrichment in various MSigDB Oncogenic Signatures sets such as KRAS, p53, PTEN, mTOR and SNF5 [KRAS.AMP.LUNG_UP.V1 _UP, P53_DN.V2_DN and PTEN_DN.V1_UP] etc. (p-value<0.05).

#### Multi-class classification

With aim to identify the features that can classify normal, early and late stage tissue samples, we have independently selected 33 CpG sites (multiclass-CpG) and 5 RNA transcripts (multiclass-RNA) using WEKA. The features are selected from 440 CpG sites and 236 RNA transcripts that are significantly differentially methylated and differentially expressed respectively (between 3 groups: cancer v/s normal and early stage v/s late stage). Here prediction models developed based on multiclass-CpG using various techniques of WEKA. Naïve Bayes model is top performer with accuracy 77.43 and 76.54 and weighted average AUROC 0.88 and 0.86 on training and independent validation dataset respectively (Table 6). The Boxplot represents the methylation pattern of multiclass-CpG features in Figure_S9A (Additional file 11). Also, we developed prediction models based on multiclass-RNA by implementing different techniques of WEKA. Naïve Bayes model achieve maximum performance with accuracy 72.73% and 72.84 and weighted average AUROC 0.81 and 0.80 on training and independent validation dataset respectively (Table 6). Boxplot represents the expression pattern of multiclass-RNA transcripts in Figure_S9B (Additional file 11). In addition, we have also developed prediction model based on 284 features selected by WEKA from all 60,483 RNA transcripts using different techniques. Model based on random forest (tree based algorithm) classify samples with accuracy 78.99% and 70.37%, weighted average AUROC 0.88 and 0.80 of training and independent dataset (Additional file 1, Table_S13). The performance of models exploiting other techniques lowers than of above mentioned models. Complete results of prediction models based on 33 CpG sites 5 and 284RNA transcripts using various techniques implementing WEKA have been shown in (Additional file 1, Table_S14).

**Table 6:**
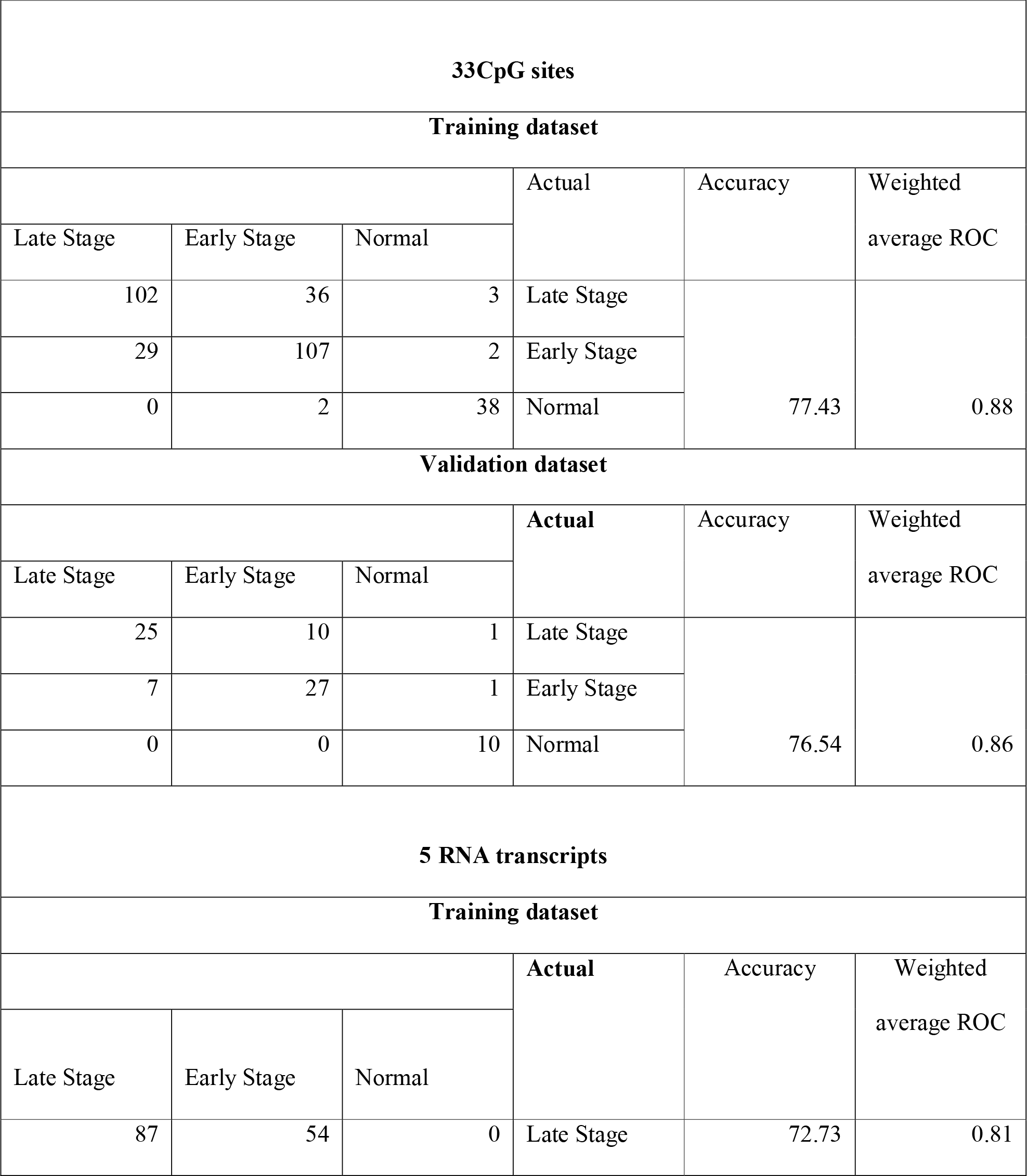
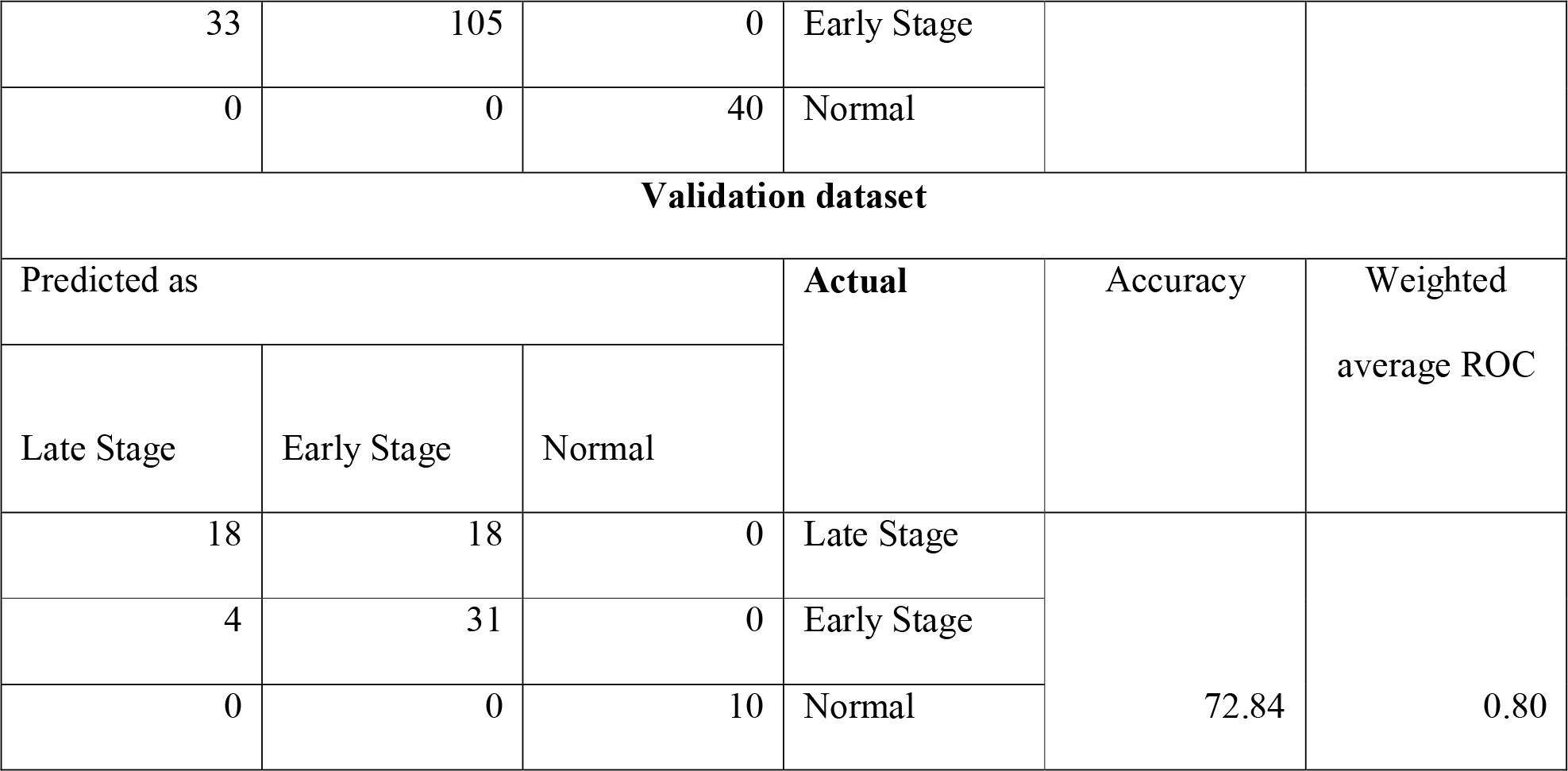
The performance Naïve Bayes model in form of confusion matrix developed for classifying normal, early and late stage samples. Model was developed using33 CpG sites (multiclass-CpG) and 5 RNA transcripts (multiclass-RNA).

| 33CpG sites |  |  |  |  |  |
| --- | --- | --- | --- | --- | --- |
| Training dataset |  |  |  |  |  |
|  |  |  | Actual | Accuracy | Weighted average ROC |
| Late Stage | Early Stage | Normal |  |  |  |
| 102 | 36 | 3 | Late Stage | 77.43 | 0.88 |
| 29 | 107 | 2 | Early Stage |  |  |
| 0 | 2 | 38 | Normal |  |  |
| Validation dataset |  |  |  |  |  |
|  |  |  | Actual | Accuracy | Weighted average ROC |
| Late Stage | Early Stage | Normal |  |  |  |
| 25 | 10 | 1 | Late Stage | 76.54 | 0.86 |
| 7 | 27 | 1 | Early Stage |  |  |
| 0 | 0 | 10 | Normal |  |  |
| 5 RNA transcripts |  |  |  |  |  |
| Training dataset |  |  |  |  |  |
|  |  |  | Actual | Accuracy | Weighted average ROC |
| Late Stage | Early Stage | Normal |  |  |  |
| 87 | 54 | 0 | Late Stage | 72.73 | 0.81 |
| 33 | 105 | 0 | Early Stage |  |  |
| 0 | 0 | 40 | Normal |  |  |
| **Validation dataset** |  |  |  |  |  |
| Predicted as |  |  | Actual | Accuracy | Weighted average ROC |
| Late Stage | Early Stage | Normal |  |  |  |
| 18 | 18 | 0 | Late Stage | 72.84 | 0.80 |
| 4 | 31 | 0 | Early Stage |  |  |
| 0 | 0 | 10 | Normal |  |  |

Enrichment analysis of multiclass-RNA is represented in (Additional file 12, Figure_S10 (A & B)). *MT1E*, *CNDP1*, *C7*, *GMNN* and *EEF1A2* are among the 5 RNA transcripts identified in multiclass classification. Enrichment analysis indicated upregulated genes such as *GMNN* and *EEF1A2* enriched in various growth promoting processes and pathways, while downregulated genes like *CNDP1* and *MT1E* enriched in negative regulators of growth and *C7* enriched in innate immunity pathways respectively.

##### Implementation of Webserver

To serve the scientific community, we developed CancerLSP (Liver cancer stage prediction), a web server that executes in silico prediction models developed in the present study for prediction of cancer status *i.e.* stage and analysis using methylation profiling and RNA expression data. Further, CancerLSP mainly has two modules; Prediction Module and Data analysis Module.

##### Prediction Module

This module permits the users to predict the disease status i.e. cancerous or normal and stage of cancer of sample using methylation beta values of CpG sites and FPKM quantification values of RNA transcripts. Here, the user required to submit FPKM values of RNA markers (RNA transcripts) and methylation beta values of marker CpG sites of every subject. The output result displays a list for patient samples and corresponding predicted status cancer or normal and early or late stage. Moreover, the user can select among the models developed from LS-CpG-WEKA, LS-RNA-WEKA LS-CpG-RNA-hybrid, LCN-5-CpG, LCN-5-CpG, multiclass-RNA and multiclass-RNA which have been identified as putative markers sets for stage progression of LIHC.

##### Data analysis Module

This module is beneficial in assessing the role of each RNA transcript and CpG site to distinguish early stage from the late stage. In addition, it provides *p*-value for every feature that indicates whether there is significant difference in methylation pattern of CpG site or RNA expression in early and late stage. Furthermore, it also provides threshold-based ROC of each feature along with mean methylation of CpG site and mean expression of that RNA transcript in the early and late stage of LIHC. This web server is available from URL http://webs.iiitd.edu.in/raghava/cancerlsp/ for scientific community.

## Conclusion and Discussion

Worldwide HCC has become a major threat owing to its high morbidity and mortality rate. Furthermore, the appropriate treatment options are affected not only by the degree of liver dysfunction but also by tumor stage[34]. Therefore, accurate stage classification and prediction of disease at early stage is crucial for patient management. Hence, the primary goal of the present study is to identify the genomic and epigenomic features that distinguish early stage from the late stage of LIHC based on the TCGA datasets retrieved from GDC data portal. Accordingly, we have made an attempt to distinguish early stage from late stage tissue samples of LIHC employing methylation profiling and RNA expression profiling data. Firstly differentially methylated CpG sites and differentially expressed transcripts identified and are ranked based on AUROC to categorize early stage and late stage tissue samples using simple threshold-based classification method.

3 CpG sites i.e. cg07132710, cg11232136 and cg03108697 of LS-CPG-AUROC signatures, are hypomethylated in early stage of LIHC are associated with *GATA2, ZNF566* and *SWAP70* respectively. Earlier studies have shown that these genes are epigenetically repressed and associated with cancer progression in different malignancies [35–38].

Various genes such as *NCAPH*, *CYP4A*, *GLYATL1*, *HILPDA*, *CANT1*, *SLC2A2*, *ALDOA*, *CNFN*, *A1BG*, *CYP2B6*, *HN1* and *ITIH1 etc.* are among the LS-RNA-AUROC signatures. Previously, different reports uncover their involvement in a variety of malignancies [39–46]. Evidently, recent studies reveal the association of *HN1* and *SLC2A2* with survival of HCC patients and *ITIH1* with progression of HCC [47–49]. Beside the protein coding genes, LINC01146 a long non coding RNA was earlier identified as differentially expressed in HCC [50,51] can classify stage samples with AUROC 0.65.

Further using state of art machine learning techniques 21 CpG (LS-CPG-WEKA) methylation sites achieved AUROC of 0.78 on validation data and 30 RNA transcripts (LS-RNA-WEKA) independently distinguish early stage from late stage tissue samples of validation dataset with 0.77 AUROC.

Among LS-CPG-WEKA signature CpG sites such as cg16657244, cg11232136, cg11023721 and cg27111890 are associated with *NOLC1, ZNF566, SMAD7* and *UBASH3A* respectively. Previously the aberrant methylation pattern of these genes has been observed in different types of cancers [38, 52, 53].

Furthermore, among the identified RNA transcripts in LS-RNA-WEKA to classify early stage and late stage of LIHC, out of 15 protein coding transcripts *DCK*, *CDCA5*, *RPP25*, *FUT11*, *NECAB1*, *FNTB*, *ZNF576*, *NETO2* and *ELOVL3* are observed as upregulated genes in late stage and are involved in cell division processes, transcriptional regulation, while genes upregulated in early stage such as *MAT1A*, *LCAT*, *CFHR3* are involved in normal functioning of liver involving lipid and lipoprotein metabolism [54] (Table_S6). Previously, various studies revealed the role of *MAT1A* in HCC [55–57]. Two of the genes in our biomarker panel *i.e. CDCA5*and *RAMP3* already identified as prognostic markers in HCC in literature [58,59]. In recent past, role of pseudogenes has been revealed in pathogenesis of cancer [60–62] including HCC (25391452, 28823960). LS-RNA-WEKA also contains 12 pseudogenes (9 processed, 2 unprocessed and 1 transcribed processed pseudogenes) including RP3-375P9.2, GAPDHP63, PTMAP2, RPS3P7, FABP5P3, HNRNPA1P37, AC018712.2 *etc.* They are significantly upregulated in late stage of HCC. One of the studies reveals the role of RP3-375P9.2 in breast cancer [63].

Models based on 21 methylation CpG sites and 30 RNA transcripts achieved reasonable performance independently for classifying early and late stage tissue samples. On combining these features, hybrid models are able to classify samples with higher performance with accuracy of nearly 78% and AUROC 0.82 on independent validation dataset.

In this study, we have also identified top 10 CpG sites and top 10 RNA transcripts as candidate signatures based on their individual performance in terms of AUROC >0.9 to classify LIHC samples and normal liver tissue samples using simple threshold based approach. *CLEC4G* and CpG site cg07274716 (associated with *PITX1*) distinguish cancer samples from normal with AUROC 0.98 and 0.97 respectively. Earlier studies have shown the association of downregulation of CLEC4G expression with the progression of HCC [64,65]. Our study corroborates this signature marker in HCC and further extends the importance of this gene as prognostic marker. Previous reports indicate that the hypermethylation of the *PITX1* correlated with tumor progression HNSCC [66] and ESCC [67]. Similarly, present study reveals the association hypermethylation of *PITX1* with progression of HCC.

Furthermore, prediction models developed using multiple features filtered by different techniques to distinguish cancer from control samples. Interestingly, our study shown that the model based on 5 features (RNA transcripts or CpG sites) performed quite similar to that of models based on much large dimension of features as indicated by the performance to discriminate cancer and normal samples with 96.47% accuracy and AUROC 0.97 on independent validation dataset. In current study, *CLEC4G*, *GDF5*, *CLEC1B*, *CLEC4M* and *FCN2* are among *LCN-5-RNA* signature that can classify cancer and normal tissue sample with high precision. Earlier, *CLEC4M* and *GDF5* have been recognized as differentially expressed gene in various other cancers [68-70], while *CLEC4G, CLEC1B* and *FCN2* were identified as genes that associated with cancer progression and prognosis in hepatocellular carcinoma [64, 71–73]. Overall this analysis emphasizes the role of these signature genes in oncogenesis. Interestingly one of the genes, *LCAT* is able to classify cancer and normal samples with AUROC 0.92, is also one of important markers to distinguish late stage samples from early stage. Thus our report suggests the role of *LCAT* as important signature marker progression of LIHC.

In addition to binary class classification, we have also developed two multiclass prediction models that can distinguish normal, early stage and late stage tissue samples based to 5 RNA transcripts sites with 72.84% accuracy with weighted AUROC of 0.81 and 33 CpG sites accuracy of 77.54% and weighted AUROC 0.86. Their enrichment analysis shows that upregulated genes enriched in various growth enhancing processes and pathways such as positives regulation of various kinase signaling, GTPase; while downregulated genes enriched in negative regulators of growth processes and innate immunity pathways. This might lay insight toward their involvement in progression of cancerous condition.

We have achieved only reasonable performance to classify early stage tissue samples from late stage tissue samples despite we tried different approaches to classify early stage tissue samples from late stage using methylation and expression data. These results suggest that stage classification is quite difficult and challenging task in comparison to cancer versus normal classification on the basis of their expression and methylation profiling. Moreover, HCC is a complex disease with various pathological bases, thus the exploration of biomarker combinations might offer more accurate and precise diagnosis or prognosis of HCC.

## Potential Implications

We anticipate this study would be beneficial to recognize these important epigenetic and genomic-based signature markers for early diagnosis of liver cancer. We have integrated all our models in the form of webserver named CancerLSP for the use of scientific community. It has mainly two modules; Prediction Module and Data analysis Module. The prediction module allows the users to predict the disease status of sample *i.e.* early or late stage of cancer and cancerous or normal using methylation beta values of CpG sites and FPKM quantification values of RNA transcripts. Further, Data analysis module is beneficial in assessing the role of each RNA transcript and CpG site to distinguish early stage from the late stage

## Methods

### Pre-processing of Data

#### Methylation Data

There are total 4,85,577 Methylation CpG site (Probe IDs associated with CpG sites) for each tissue sample. The methylation score for every CpG site was defined in terms of beta value. All those CpG sites approximately 23-25% excluded from the study for which beta value is missing among any of samples using in-house bash script. Hence CpG sites number reduced to 3,74,292 for staging analysis and 3,69,221 for Cancer v/s Normal analysis.

#### Normalization of RNA Expression

The expression values of RNA transcripts from GDC portal were obtained in term of FPKM (fragments per kilobase of transcript per million mapped reads). There is a wide range of variation in FPKM values, thus we transformed values using log2 after addition of 1.0 as a constant number to each of FPKM value. Further those features have been removed that have low variance using *caret* package in R, followed by z-score normalization of data. Thus, for each mRNA log_2_-transformed FPKM values were centered and scaled by employing *caret* package in R. Following equations were used for computing the transformation and normalization:

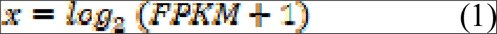

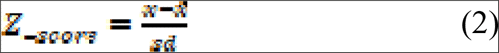

Where *Z__score_* is the normalized score, *x* is the log-transformed expression, x̄ is the mean of expression and *sd* is the standard deviation of expression.

#### Identification of differentially methylated CpG sites and expressed RNA transcripts

In order to identify differentially CpG methylated sites, first we computed the mean methylation score for each CpG site in early and late stage samples. Secondly, we computed whether difference in the mean of methylation score in early and late stage samples is statistically significant or not using *t-test*. Here, we implemented Welch t-test using in-house Rscript and bash script. Similarly, we have identified differentially expressed RNA transcripts in early and late stage samples from total of 60,483 RNA transcripts.

#### Feature Selection Techniques

One of challenges in developing prediction model is to identify important features from the large dimension of features. In this study, we used number of techniques for feature selection. Firstly, we used Area under Receiver operating characteristic (AUROC) based feature selection technique in which we developed single feature based models for discriminating early and late stage samples. Single feature based models are also called threshold based models in which feature having a score above threshold is assigned to early stage if it is upregulated in early stage [74]. We compute performance of each model based on a given feature and identify features having highest performance in term of AUROC.Secondly, we perform feature selection using two different algorithms like employing attribute evaluator named, ‘SymmetricalUncertAttributeSetEval’ with search method of ‘FCBFSearch’ of WEKA software package and sklearn.feature_selection. F-ANOVA method using Scikit package. We also filtered CpG sites and RNA transcripts as features that can discriminate early stage from late stage cancer samples using attribute evaluator named, ‘SymmetricalUncertAttributeSetEval’ with search method of ‘FCBFSearch’ of WEKA software package. The FCBF (Fast Correlation-Based Feature) algorithm employed correlation to identify relevant features in high-dimensional datasets in small feature space [75].

#### Implementation of machine learning techniques

Primarily we have developed the Support Vector Machine (SVM) based prediction models using the package SVM ^*light*[76]^ and WEKA [75]. In present study RBF (radial basis function) kernel employed to optimize various parameters to get best performance on training dataset.Furthermore, some of commonly used classifiers were also used for developing prediction models. These classifiers include Random forests, SMO, Naive Bayes, J48 were implemented exploiting WEKA software.

#### Performance Evaluation of Models

In current study, we used both internal and external validation technique to evaluate the performance of models. Main dataset has been divided in two sets called training dataset and validation dataset in ratio of 80:20. The training set is used for developing model and for performing internal validation; whereas, validation dataset is used to perform external or independent validation. In this article, we used ten-fold cross validation technique where training dataset is randomly split into ten sets; of which nine out of ten sets were used as training sets and the remaining tenth set as testing dataset. This process is repeated ten times in such a way that each set is exploited once for testing. The final performance of model is the mean performance of all the ten sets. In order to avoid optimization of parameters in case of ten-fold cross-validation, we also implement external validation. In case of external validation, we evaluate our model on an independent or external dataset not used for training. In this study, we used 20% of the main dataset for validation or independent testing and remaining 80% dataset for training.

In order to measure performance of models, we used standard parameters, commonly used to measure performance of classification models. Both threshold-dependent and threshold-independent parameters were employed to measure performance. In case of threshold-dependent parameters, we measure sensitivity, specificity, accuracy and Matthew’s correlation coefficient (MCC) using following equations.

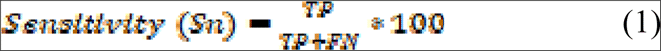

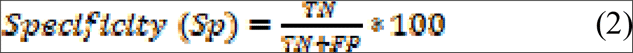

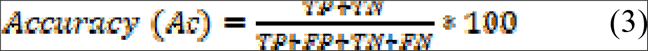

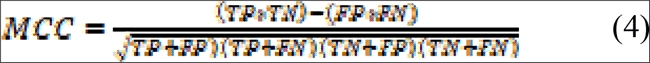

where, FP, FN, TP and TN are false positive, false negative true positive and true negative predictions respectively.

While, for threshold-independent measures, we used standard parameter Area Under the Receiver Operating Characteristic curve (AUROC) or commonly known as Area Under (AUC) or Receiver Operating Characteristic curve (ROC). The AUROC curve is generated by plotting sensitivity or true positive rate against the false positive rate (1-specificity) at various thresholds. Finally, the area under AUROC curve calculated to compute a single parameter called AUROC.

#### Functional annotation or enrichment analysis of genes

In order to discern the biological relevance of the signature genes or the genes associated with signature CpG sites, enrichment analysis performed using Enrichr [77,78]. Enrichr applies Fisher exact test to identify enrichment score. It also provides Z-score which is derived by applying correction on a Fisher Exact test.

## Contributions

H. K. and S.B. collected the data and created the datasets, developed classification programs, implemented algorithms, created the back-end server and front-end user interface. H.K., S.B. and G.P.S.R. analyzed the results. H.K. and S.B. wrote the manuscript. G.P.S.R. conceived and coordinated the project, helped in the interpretation and analysis of data, refined the drafted manuscript and gave complete supervision to the project. All of the authors read and approved the final manuscript.

## Competing interests

The authors declare no competing financial interests.

## Acknowledgement

The authors acknowledge funding agencies J. C. Bose National Fellowship (DST). HK and SB are thankful to CSIR and ICMR for providing fellowships.

## Abbreviations

MCC: Matthews correlation coefficient; AUROC: Area under Receiver Operating Characteristics; FDR: False Discovery Rate

